# Statistical approach for …*a test chemical is considered to be positive*… in regulatory toxicology: Trend and pairwise tests

**DOI:** 10.1101/858571

**Authors:** Ludwig A. Hothorn

## Abstract

In regulatory toxicology an outcome is claimed positive when both a trend is significant and any pairwise test against control. Two statistical approaches are proposed: a joint Dunnett and Williams test (assuming the dose as a qualitative factor) and a joint test of the Tukey regression test and Dunnett test (assuming the dose as a quantitative covariate). Related R software is available.

## 1 The problem

In regulatory toxicology, different bioassays are routinely evaluated (commonly on the basis of multiple endpoints) into positive or negative outcomes. In most cases, the criteria are not clearly formulated from a statistical point of view. However, there are guidelines with explicit criteria, such as OECD No. 487: … *a test chemical is considered to be positive if*…: *1)At least one of the treatment groups exhibits a statistically significant increase in the frequency of micronucleated immature erythrocytes compared with the concurrent negative control, 2) This increase is dose-related at least at one sampling time when evaluated with an appropriate trend test, and 3) c) Any of these results are outside the distribution of the historical negative control data.…* Although criterion 3 is also challenging from a statistical point of view, a statistical approach is derived below for the first two criteria: **pairwise tests and trend test**.

Without limitation of generalizability, only tests for normal distributed and homoscedastic errors are represented here, in order to keep the calculation simple.

## 2 Methods

Various criteria must be observed. First, a trend test should be sensitive to as many forms of dose-response dependency as possible. Here the Williams test is selected for dose modeling as a qualitative factor and the Tukey test for dose modeling as a quantitative covariate. Secondly, pairwise tests are only considered as comparisons against control (i.e. not between doses comparisons) in terms of the widespread Dunnett test [5] (which controls familywise error rate). Sometimes independent t-tests are simply used for this purpose. However, these only control the comparisonwise error rate, use only pairwise df and error estimates. Thirdly, only one-sided tests are used because a test is difficult to motivate which is either to formulate for an increase or a decrease, but this for monotonous dependencies. The combination of trend test and Dunnett-test takes place on the level of linear models [8]. On the one hand, by the simultaneous consideration of Williams and Dunnett contrasts [9], on the other hand by the simultaneous testing of Tukey [10] and Dunnett test [7]. Simultaneous means the use of the common joint distribution of all tests in the sense of a maximum test.

This shows a fundamental contradiction in this approach. On the one hand, claims are possible with regard to trend and any pair comparison in the sense of the guideline, on the other hand, the false negative decision rate increases. A simulation study shows that the balancing of interests: extended claim - despite of increased f-rate is acceptable. With the help of two examples the advantages and disadvantages of the approach are shown, the data and the R-code are made available, so that a recalculation of own data should be possible for toxicologists.

The colloquial “and” is statistically translated into “or” by a union-intersection hypothesis. A trend exists if at least one local trend alternative is significant, a pairwise comparison if at least one of the comparisons is significant. Of course, different patterns may emerge from significant local alternatives until all are significant. Conversely, if none of the local comparisons are in the alternative, there is no trend that is only a paired difference (see the examples below).

## 3 Data examples

### 3.1 Number of revertants in Ames assay

The raw data of an Ames assay using TA98 is available in the R-library(dispmod) taken from [4]. A clear downturn effect can be see above a dose of 333.

**Figure 1:**
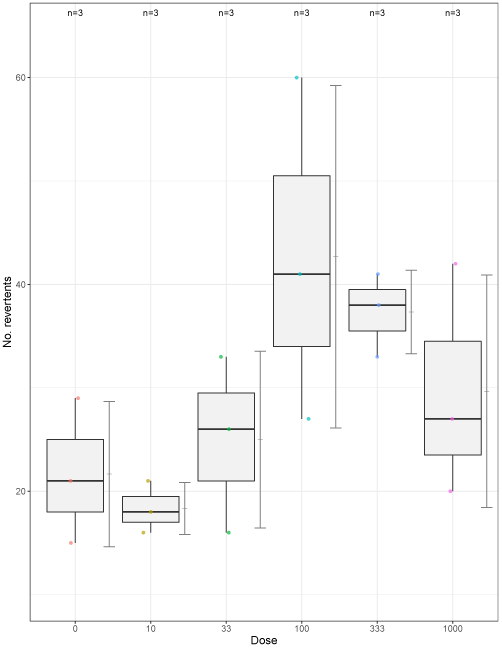
Box-plot Ames assay.

The Tukey-and-Dunnett approach is used by means of the library(tukeytrend) and the function *tukeytrendfit*, where a linear model for the log-transformed number of revertants (y) is used as an object (modTA). The object ex2 contains the test statistics and the multiplicity-adjusted p-value (see Table).

~~~
library(dispmod)
data(salmonellaTA98)
modTA<-lm(log(y)∼dose, data=salmonellaTA98)
library(tukeytrend)
EX2 <-tukeytrendfit(modTA, dose=“x”,
 scaling=c(“ari”, “ord”, “arilog”, “treat”), ctype=“Dunnett”)
ex2<-summary(asglht(EX2, alternative=“greater”))
~~~

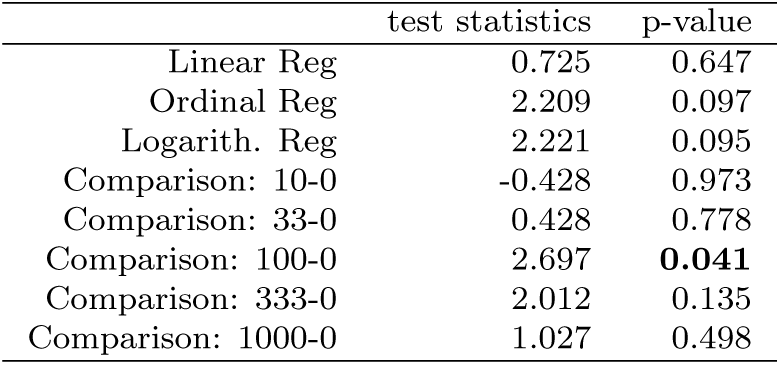

Both linear trend test and Tukey trend test alone argue for ‘no trend’, where the joint approach reveal a significant increase in dose 100.

### 3.2 Crude tumor rates in a long-term carcinogenicity study

The crude incidence of hepatoblastoma in male mice (1,1,16,5) were reported after treatment of 0 3 15 50 mg/kg pentabrominated diphenyl ether [6] (*n*_*i*_ = 50). Assuming dose as a qualitative factor, the proportions were modeled in the generalized linear model with the logit link function and small sample add-2 sample adjustment [2].

~~~
library(multcomp)
ta<-data.frame(
 dose = c(0, 3, 15, 50),
 tumor = c(1,1,16,5),
 mice = c(50,50,50,50))
ta$Dose<-as.factor(ta$dose) # dose as factor
modAC<-glm(cbind(tumor+.5,(mice-tumor)+.5)∼Dose, family=binomial, data=ta)
nn<-table(ta$Dose)
matC<-contrMat(nn, type=“Dunnett”) # Dunnett contrasts
matW<-contrMat(nn, type = “Williams”) # Williams contrasts
matCW<-rbind(matC,matW) # DunWil contrasts
plot(glht(modAC, linfct = mcp(Dose =matCW))) # joint test
~~~

The simultaneous confidence limits (on the log-odds ratio scales) for the six individual comparisons: i) Dunnett-type vers. control, and ii) Williams-type C1 for 0 − 50, C2 for 0 − (50 + 15)*/*2, C3 for 0 − (50 + 15 + 3)*/*3 reveal no trend, but a significant increase of the tumor rate at dose 15.

**Figure 2:**
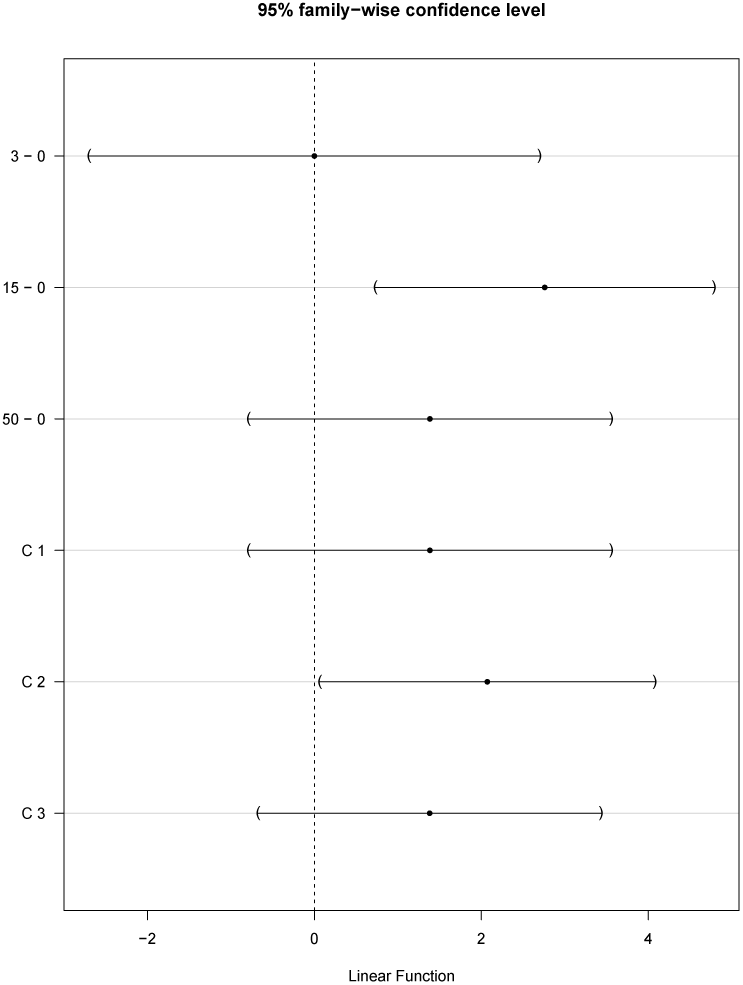
Confidence intervals for Dunnett-Williams contrasts.

## 4 Simulation study

In a simulation study for a common balanced design [*NC, D*_1_, *D*_2_, *D*_3_] with homoscedastic, normal distributed errors with small sample sizes (main *n*_*i*_ = 10 and down to *n*_*i*_ = 4 the false positive rate (under the global null hypothesis of equal expected values (means) and the false negative rate under various monotonic and non-monotonic dose-response relationships. Seven tests were compared: i) the standard Dunnett-test (Dun), the Williams test formulated as multiple contrast [3] (Wil), the joint Dunnett and Williams test (DunWil), the Tukey trend tests (Tukey), the joint Tukey and Dunnett test (TukDun), the linear regression test (Lin) and the joint linear Regression and Dunnett test (LinDu), see the following Table:

**Table 1:** False negative decision rates.

| Alternat | Shape | Dun | Will | DunWil | Tukey | TukDu | Lin | LinDu |
| --- | --- | --- | --- | --- | --- | --- | --- | --- |
| $H_0$ | $\mu, \mu, \mu, \mu$ | 0.951 | 0.948 | 0.952 | 0.953 | 0.952 | 0.948 | 0.951 |
| Mono | $\mu, \mu, \mu, \mu + \delta$ | 0.112 | 0.081 | 0.117 | 0.035 | 0.062 | 0.036 | 0.058 |
| Linear | $\mu, \mu + \delta/3, \mu + \delta/2, \mu + \delta$ | 0.095 | 0.061 | 0.087 | 0.038 | 0.070 | 0.037 | 0.078 |
| Mono | $\mu, \mu, \mu + \delta, \mu + \delta$ | 0.040 | 0.027 | 0.036 | 0.017 | 0.019 | 0.096 | 0.032 |
| Plateau | $\mu, \mu + \delta, \mu + \delta, \mu + \delta$ | 0.022 | 0.010 | 0.018 | 0.120 | 0.027 | 0.515 | 0.026 |
| Non-mo | $\mu, \mu, \mu + \delta, \mu$ | 0.115 | 0.498 | 0.123 | 0.712 | 0.141 | 0.979 | 0.133 |
| Non-mo | $\mu, \mu + \delta, \mu, \mu$ | 0.117 | 0.699 | 0.121 | 0.999 | 0.137 | 0.999 | 0.130 |
| Non-mo | $\mu, \mu + \delta/3, \mu + \delta, \mu$ | 0.120 | 0.476 | 0.123 | 0.903 | 0.142 | 0.993 | 0.132 |
| Non-mo | $\mu, \mu, \mu + \delta, \mu + \delta/3$ | 0.109 | 0.262 | 0.111 | 0.296 | 0.108 | 0.760 | 0.119 |
| Non-mo | $\mu, \mu + \delta/3, \mu + \delta, \mu + \delta/3$ | 0.101 | 0.243 | 0.103 | 0.491 | 0.112 | 0.868 | 0.111 |
| Non-mo | $\mu, \mu + \delta/3, \mu + \delta, \mu + \delta/2$ | 0.090 | 0.091 | 0.082 | 0.166 | 0.079 | 0.467 | 0.093 |
| Lin $n_i = 11$ | $\mu, \mu + \delta/3, \mu + \delta/2, \mu + \delta$ | 0.078 | 0.050 | 0.070 | 0.039 | 0.058 | 0.067 | 0.059 |
| Lin $n_i = 10$ | $\mu, \mu + \delta/3, \mu + \delta/2, \mu + \delta$ | 0.095 | 0.061 | 0.087 | 0.038 | 0.070 | 0.037 | 0.078 |
| Lin $n_i = 9$ | $\mu, \mu + \delta/3, \mu + \delta/2, \mu + \delta$ | 0.126 | 0.083 | 0.121 | 0.055 | 0.088 | 0.108 | 0.097 |
| Lin $n_i = 8$ | $\mu, \mu + \delta/3, \mu + \delta/2, \mu + \delta$ | 0.151 | 0.111 | 0.150 | 0.083 | 0.118 | 0.139 | 0.130 |
| Lin $n_i = 7$ | $\mu, \mu + \delta/3, \mu + \delta/2, \mu + \delta$ | 0.216 | 0.174 | 0.207 | 0.127 | 0.172 | 0.215 | 0.188 |
| Lin $n_i = 6$ | $\mu, \mu + \delta/3, \mu + \delta/2, \mu + \delta$ | 0.291 | 0.215 | 0.286 | 0.179 | 0.240 | 0.290 | 0.260 |
| Lin $n_i = 5$ | $\mu, \mu + \delta/3, \mu + \delta/2, \mu + \delta$ | 0.363 | 0.275 | 0.349 | 0.234 | 0.303 | 0.341 | 0.322 |
| Lin $n_i = 4$ | $\mu, \mu + \delta/3, \mu + \delta/2, \mu + \delta$ | 0.470 | 0.379 | 0.464 | 0.321 | 0.412 | 0.449 | 0.435 |

As expected, the regression test has the lowest f-rate for a linear alternative. As expected, the regression test has the lowest f-rate for a linear alternative. For many other nearly linear alternatives, the Tukey test has the lowest f-rate. For plateau alternatives, the Williams test shows the lowest f+ rates (while the regression test there shows alarmingly high f-rates, which disqualifies it as a routine approach in toxicology). As expected, the f-rates of the both regression tests are high for non-monotonous alternatives, much lower for the Williams test. The three combination tests with the additional Dunnett test show a balanced f-behavior. Their f-rate is more or less increased compared to the standard test with exactly monotonous alternatives - however tolerable due to the considerable robustness gain. This difference is about the same over wide ranges of the f-rate, represented by designs with decreasing sample sizes *n*_*i*_ = 11…*n*_*i*_ = 4 (common in toxicology).

It can be assumed that this behavior also applies to other endpoint types in GLM. Therefore, the recommendations of the CA-trend tests for crude or poly-3-adjusted proportions should be reconsidered by the US-NTP [1].

## 5 Conclusion

Two statistically consistent approaches for trend and pairwise comparisons were derived and their properties characterized: Dunnett and Williams test and Tukey and Dunnett test. A relatively simple R-code is available for both. These approaches can be generalized in GLM to other endpoint types such as proportions, counts and survival functions. Thus, this can be recommended for routine evaluation in regulatory toxicology.

Extensions for selected real data conditions such as variance heterogeneity, over-dispersion or values below the detection limit are currently being processed.

